## Supporting Figures for "Escape problem of magnetotactic bacteria - physiological magnetic field strengths help magnetotactic bacteria navigate in simulated sediments"

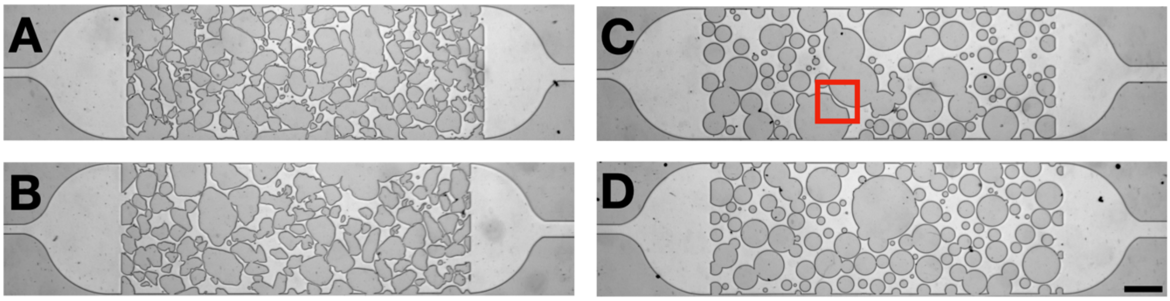


**Figure S1:** Two microfluidic channels with irregularly shaped (A,C) obstacles and the corresponding channels with rounded obstacles (B,D). Note that in (C), a narrow channel has been cut out of the big obstacle in the center (indicated by the red square), as it would block the channel completely. The scale bar is 150 μm.

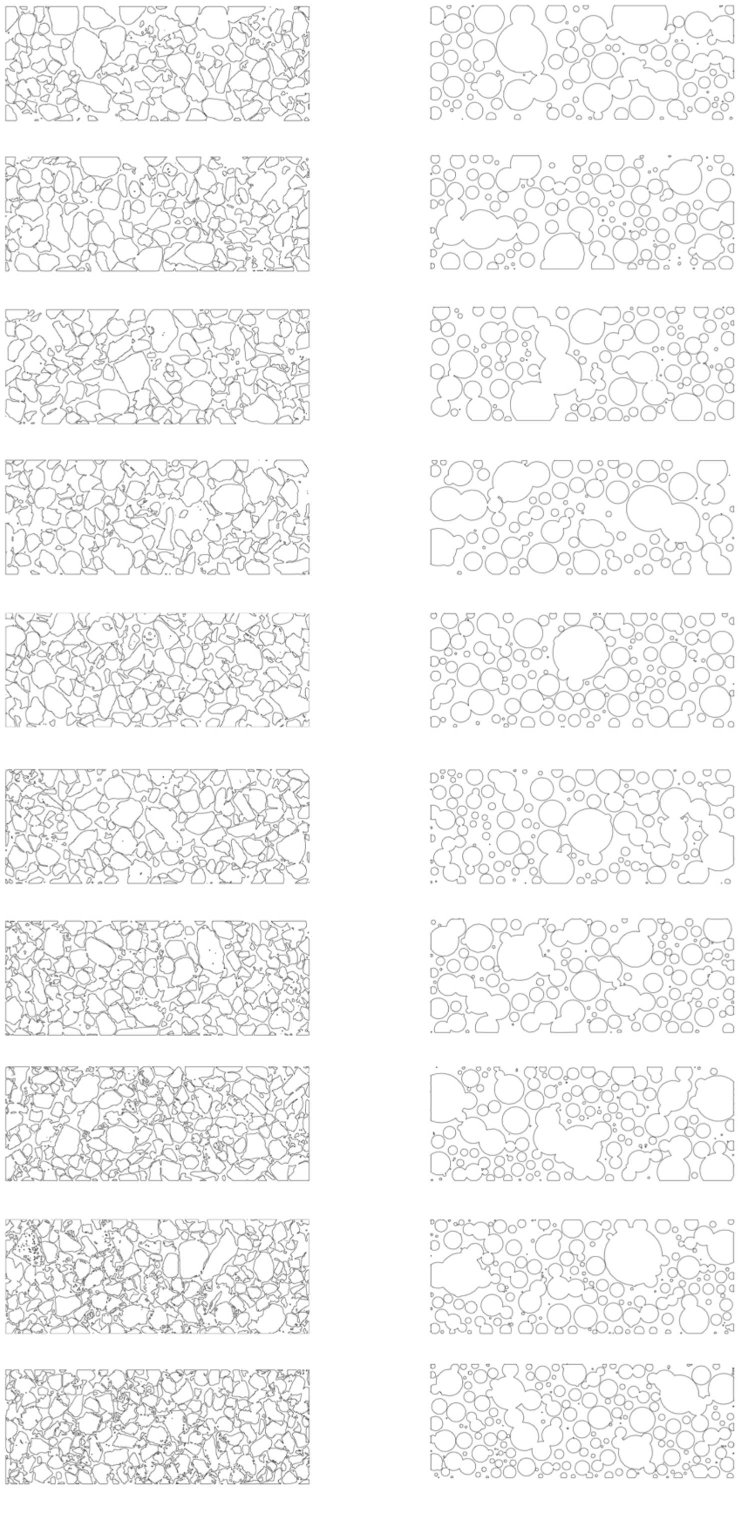

**Figure S2:** Masks for all obstacle arrays used in this study. Left column: irregular shaped obstacles based on 2d μCT images of sand samples, right column: rounded obstacle.

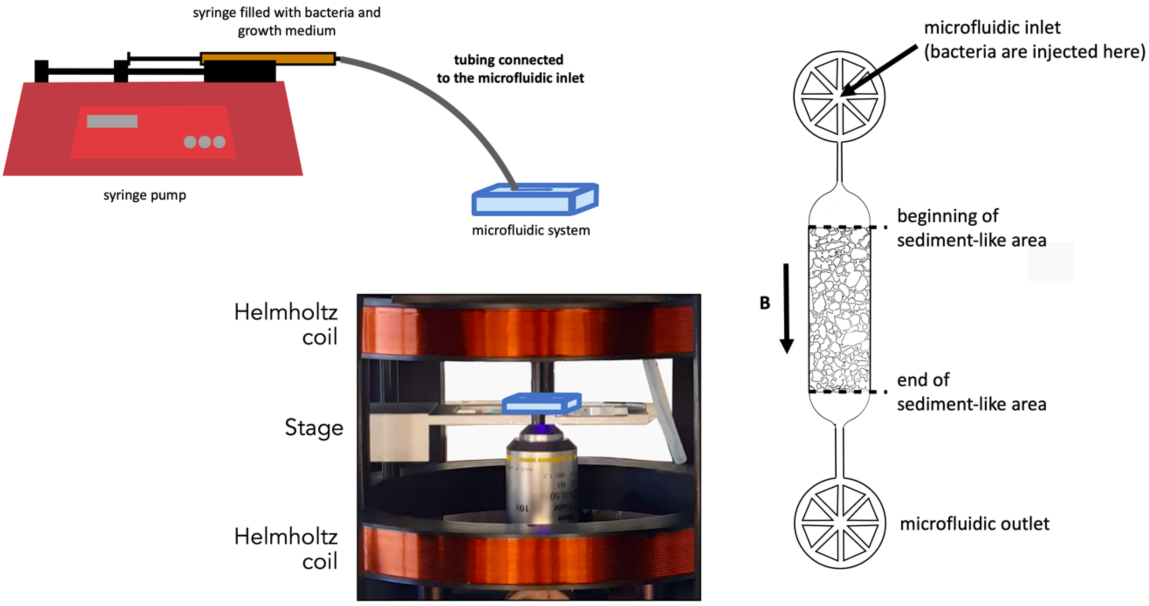


**Figure S3:** Sketch of the setup showing the microfluidic system (blue), the inlet connecting to a syringe with bacteria and growth medium, one of the three pairs of Helmholtz coils to control the magnetic field, and a detailed view of the microfluidic system.

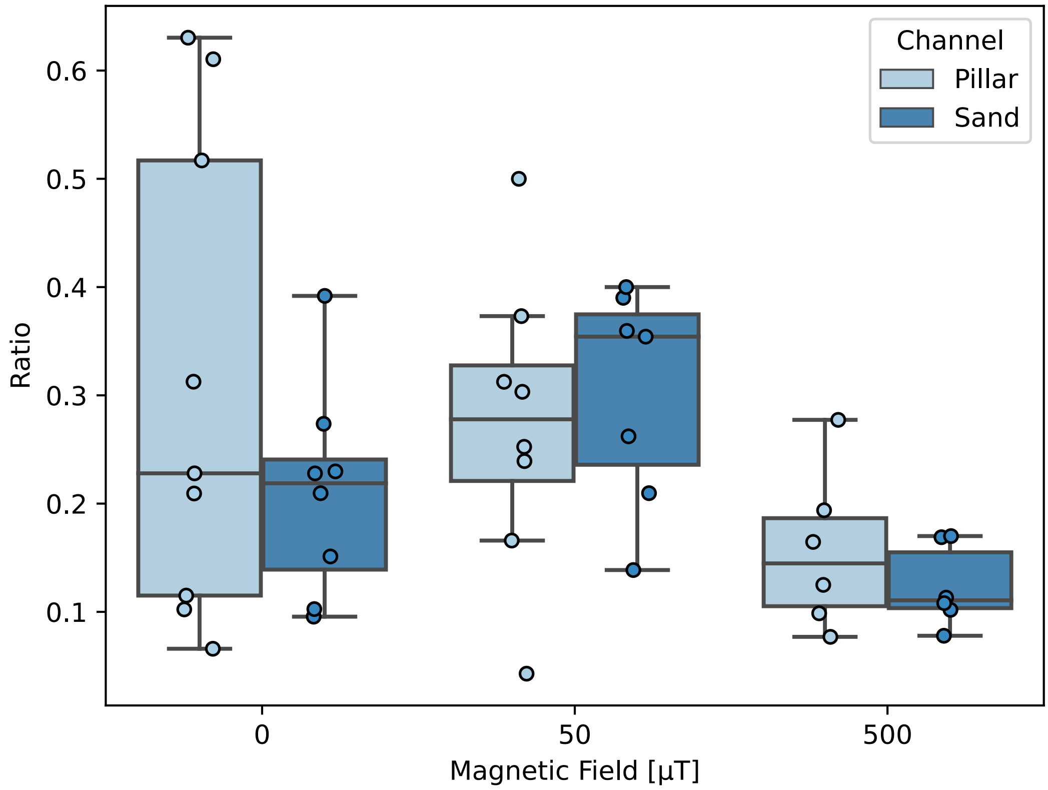


**Figure S4:** Comparison of bacterial throughput as a function of the magnetic field (0, 50, 500 μT) in channels with irregular (sand-like) obstacles and in channels with rounded (pillar-like) obstacles.

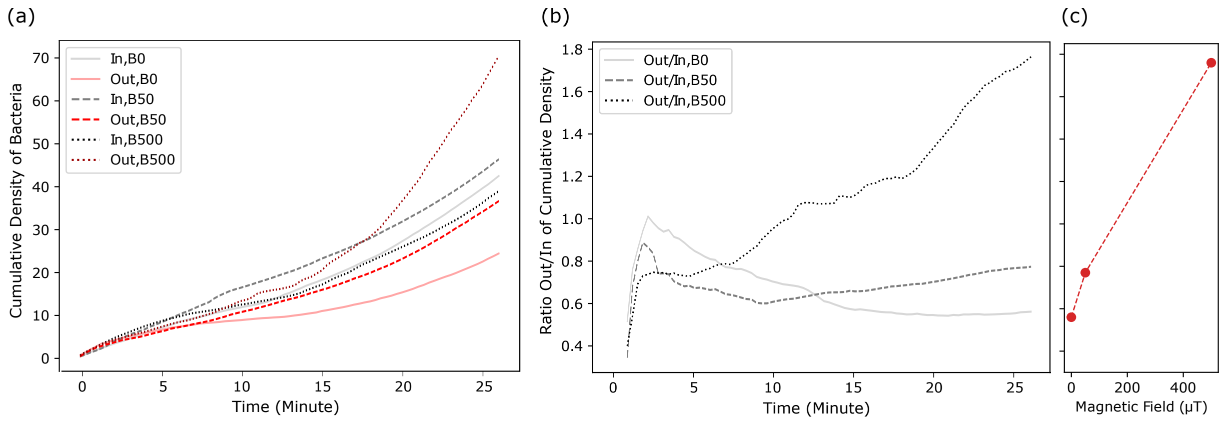


**Figure S5:** Bacterial throughput in a channel without obstacles: a) cumulative density of bacteria in the IN and Out regions of the channel. B) ratio of the cumulative density in the IN and Out regions, both as functions of time and for different magnetic field strengths. C) Dependence of the latter ratio (taken after 30 min) on the magnetic field strength. In contrast to channels with obstacles (Figure 3), a monotonic dependence is observed.

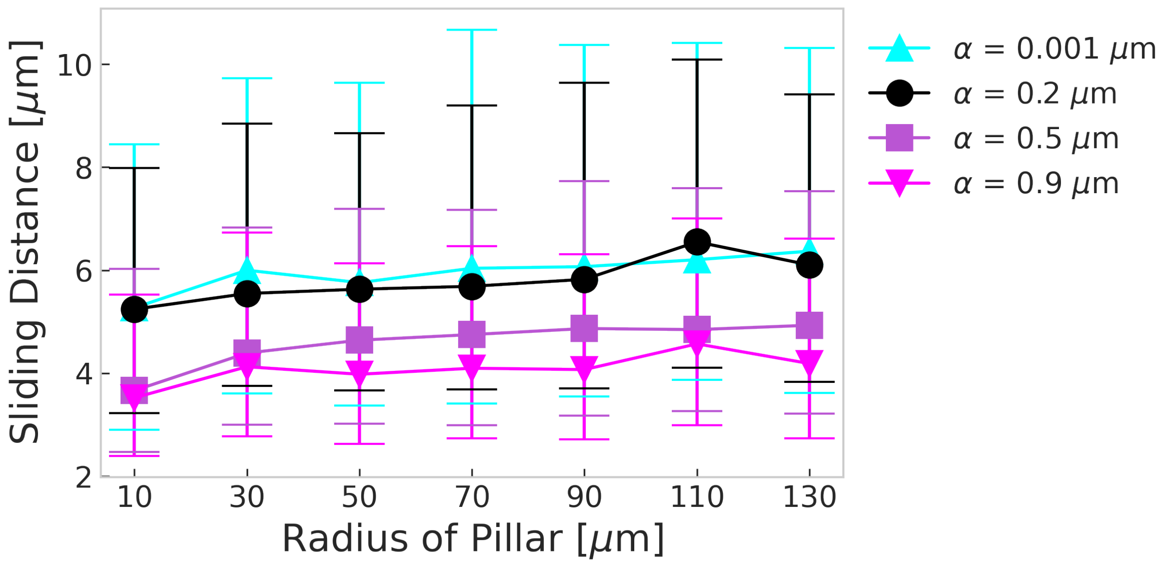


**Figure S6:** Sliding distance as a function of the pillar radius as obtained from simulations for different wall torque parameters α. The presented values show the median, error bars indicate 25th and 75th percentile.  The sliding distances were extracted from obstacle channel simulation data, where pillar radii were binned into 20 μm bin sizes for statistical analysis.

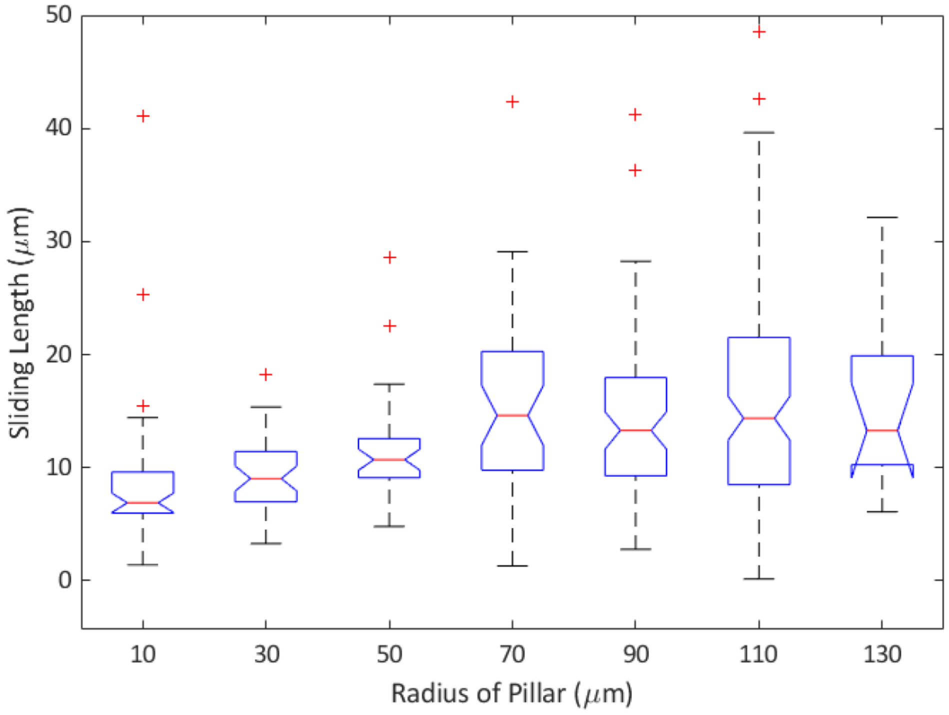


**Figure S7:** Dependence of the sliding distance on the curvature of the pillar as obtained from the tracking bacteria in obstacle channels with cylindrical obstacles.
